# Substrate stiffness modulates the emergence and magnitude of senescence phenotypes in dermal fibroblasts

**DOI:** 10.1101/2024.02.06.579151

**Authors:** Bartholomew Starich, Fan Yang, Derin Tanrioven, Heng-Chung Kung, Joanne Baek, Praful R. Nair, Pratik Kamat, Nico Macaluso, Joon Eoh, Kyu Sang Han, Luo Gu, Jeremy Walston, Sean Sun, Pei-Hsun Wu, Denis Wirtz, Jude M. Phillip

## Abstract

Cellular senescence is a major driver of aging and disease. Here we show that substrate stiffness modulates the emergence and magnitude of senescence phenotypes after exposure to senescence inducers. Using a primary dermal fibroblast model, we show that decreased substrate stiffness accelerates senescence-associated cell-cycle arrest and regulates the expression of conventional protein-based biomarkers of senescence. We found that the expression of these senescence biomarkers, namely p21^WAF1/CIP1^ and p16^INK4a^ are mechanosensitive and are in-part regulated by myosin contractility through focal adhesion kinase (FAK)-ROCK signaling. Interestingly, at the protein level senescence-induced dermal fibroblasts on soft substrates (0.5 kPa) do not express p21^WAF1/CIP1^ and p16^INK4a^ at comparable levels to induced cells on stiff substrates (4GPa). However, cells express *CDKN1a, CDKN2a,* and *IL6* at the RNA level across both stiff and soft substrates. Moreover, when cells are transferred from soft to stiff substrates, senescent cells recover an elevated expression of p21^WAF1/CIP1^ and p16^INK4a^ at levels comparable to senescence cells on stiff substrates, pointing to a mechanosensitive regulation of the senescence phenotype. Together, our results indicate that the emergent senescence phenotype depends critically on the local mechanical environments of cells and that senescent cells actively respond to changing mechanical cues.

## INTRODUCTION

Cellular senescence is a major driver of aging and is associated with a wide range of pathologies in humans^1,2^. Triggered by both intrinsic and extrinsic factors, including genotoxic, oncogenic, and replicative stresses; senescence is a phenotype characterized by proliferation arrest, resistance to apoptosis, and a pro-inflammatory secretory profile (senescence associated secretory phenotype (SASP))^3,4^. Conventionally, cellular senescence is measured based on the accumulation of β-galactosidase (SA-β-gal), upregulation of cell-cycle checkpoint pathways p16^INK4a^, p21^WAF1/CIP1^, and p53, with a shift towards larger nuclear and cellular morphologies that are observed primarily *in vitro*^3–5^. Recent findings from animal models and clinical trials in humans have shown that the depletion of senescent cells mitigates various aging-related dysfunctions and influences tissue regeneration and repair^6,7^. However, the role of extracellular mechanical cues in dictating senescence phenotypes remain poorly understood.

Senescent cells accumulate within aging tissues and accompany significant restructuring and remodeling of the tissue architecture. For instance, aging skin undergoes a thinning of the dermis and heterogeneous remodeling of the extracellular matrix (ECM)^8,9^. At the cellular scale, aged fibroblasts exhibit decreased motility^10^, a higher propensity for nuclear deformations^11^, with decreased ECM remodeling and increased deposition of collagens^8^. Together, these changes contribute to a mechanically stiffer and more fibrous extracellular microenvironments, which are associated with pathogenesis and disease^8,12^. Within the human body, healthy and diseased tissue exhibit tissue mechanics (*i.e.,* stiffness) that span several orders of magnitude ranging from bone with a Young’s modulus of ∼1-20GPa to adipose tissues having a Young’s modulus of ∼0.5-1kPa^13,14^. Within each of these mechanically tuned microenvironments, the capacity for cells to sense and transduce mechanical signals (termed mechanosensation and mechanotransduction) modulates their functions and responses to perturbations, including the capacity and rate of cell proliferation and differentiation ^14–19^. Together, this mechano-modulation of cell function provides a clear rationale to study how extracellular mechanical cues influence the emergence and onset of senescence phenotypes.

In this study, we evaluate the development of senescence in cells on substrates of defined mechanical stiffnesses. Using primary dermal fibroblasts cultured on collagen-I functionalized polyacrylamide hydrogels of various elastic moduli, we establish how senescent fibroblasts respond to changes based on the mechanical properties of their substrates, while maintaining the biochemical (ligand) presentation to the cells. We found that following senescence induction, fibroblasts are significantly more likely to enter a non-proliferative senescence state on mechanically soft substrates compared to stiff substrates (0.5kPa *vs*. 4GPa). Senescent cells did not display the expected elevated levels of p16^INK4a^ and p21^WAF1/CIP1^ at the protein level when seeded on soft substrates. The expressions of p16^INK4a^ and p21^WAF1/CIP1^ are mechanosensitive and regulated via myosin contractility through focal adhesion kinase (FAK)-ROCK signaling. Furthermore, this mechanosensitive regulation of p21^WAF1/CIP1^ is initiated at the RNA level based on the expression of senescence biomarker genes, although not manifested at the protein level on soft substrates. Our results show that the development of senescence phenotypes depend critically on the mechanics of the cellular microenvironment and that the relative expression of senescence-associated biomarkers vary with matrix stiffness.

## RESULTS

### Substrate stiffness regulates proliferation arrest during senescence induction

To evaluate the effects of substrate stiffness on senescence, fibroblasts were seeded at low density (1500 cells/cm^2^) onto collagen-I coated well plates and senescence was induced by exposing cells to 50µM of chemotherapeutic drug bleomycin for 4 h (**Figure 1a**) ^20–22^. After exposure, cells were washed 3 times with PBS to remove residual drug. As an initial measure of the senescence phenotype, we defined senescence as proliferative arrest of cells after exposure to senescence inducer bleomycin^23^. To confirm the senescence phenotype, we also evaluated the expression of conventional protein-based biomarkers which included the accumulation of senescence-associated β-Galactosidase (SA-β-Gal), stable proliferation arrest as assessed by 5-ethynyl 2’-deoxyuridine (EdU), and the upregulation of G1 checkpoint cyclin dependent kinase inhibitors (CDKi), p16^INK4a^ and p21^WAF1/CIP1^ (**Figure 1b**, **Supplemental Figure 1b, Figure 3**). Results confirmed that drug concentrations and exposure times used induced senescence on 4GPa plastic substrates at levels previously reported in the literature^4,5,22^.

**Figure 1.**
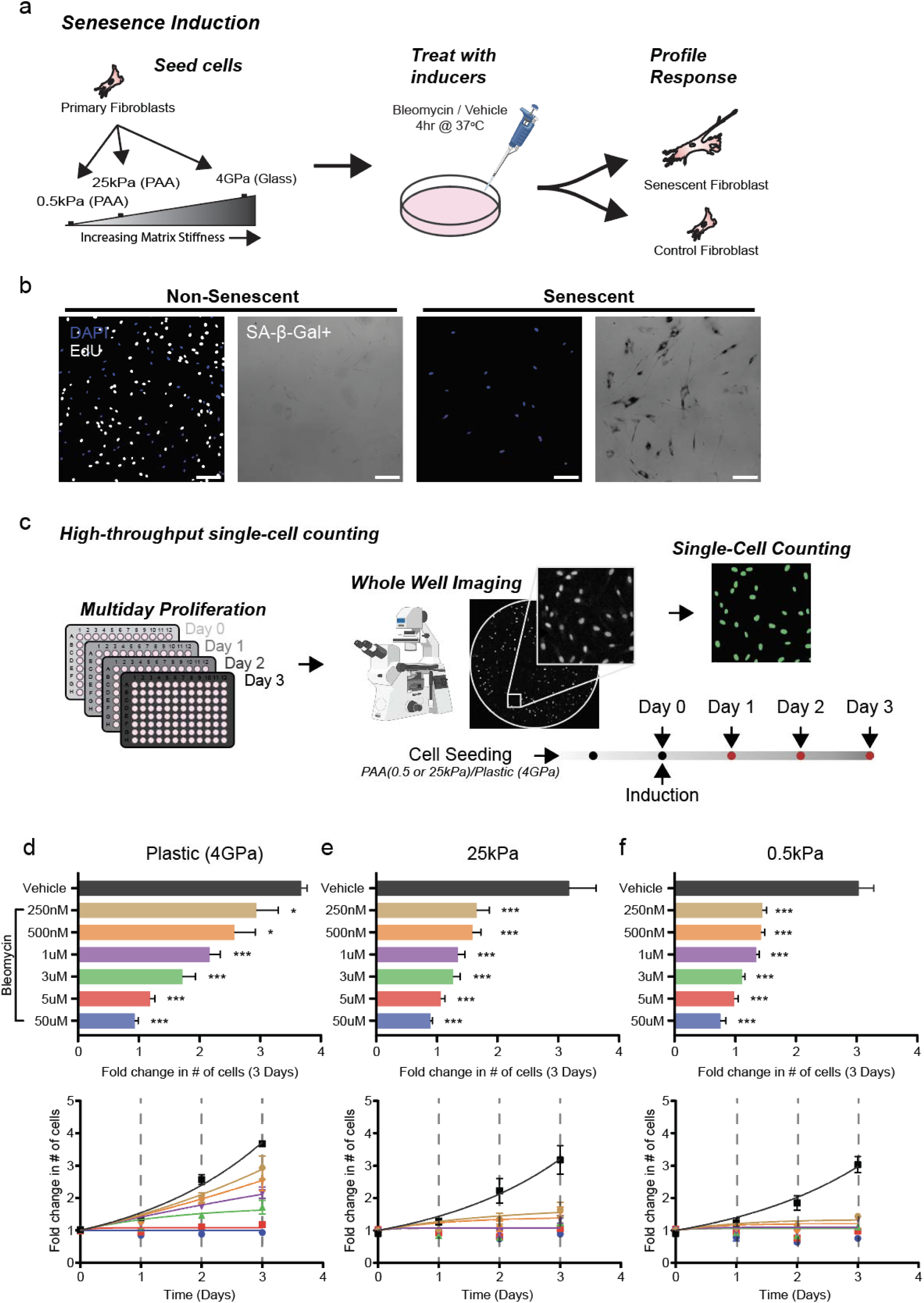
Substrate stiffness regulates proliferation arrest during senescence induction. (**a**) Drug induction workflow for biomechanical studies of cellular senescence. (**b**) GT22 Fibroblasts were seeded on collagen coated plastic substrates and treated with either bleomycin or vehicle control. Fibroblasts were stained and imaged at 10X with either fluorescent EdU or colorimetric SA-β-Galactosidase confirming the senescence phenotype, scalebar 200µm. (**c**) High-throughput single-cell counting workflow for semi-au omated whole well cell counting of GT22 nuclei over multi-day timeline. Cell counts were normalized to an initial day zero value and plotted as a function of time over the 3-day experiment. (**d-f**) Bleomycin 3-day endpoint bar plots and corresponding dose response curves showing fold change in number of GT22 fibroblasts vs time for cells subjected to Vehicle, 250nM, 500nM, 1µM, 3µM, 5µM, 50µM: (**d**) Plastic (4GPa), (**e**) 25 kPa, and (**f**) 0.5 kPa matrices. Error bars represent SEM of biological variation, statistics evaluated via one-way ANOVA, with Tukey multiple comparison test to vehicle control; data presented are from 3 biological replicates, with each having 3-5 technical replicates. (***P <0.001, **P < 0.01, *P < 0.05).

The mechanical properties of tissues are spatially heterogenous, yet the role of matrix mechanics in the emergence of senescence remains largely unclear. To evaluate the impact of matrix stiffness on initiation of senescence programs, we measured cell proliferation at three different physiologically relevant stiffnesses: 0.5 kPa polyacrylamide gels (“soft” PAA), 25 kPa PAA (“stiff”), and 4 GPa tissue culture plastic (“plastic”), corresponding to the stiffnesses comparable to that of adipose, dermal and bone tissues, respectively^13,14^. All substrates were coated with type-I collagen to maintain comparable extracellular matrix (ECM) presentation. Here, we define a functional readout of senescence as the halt in proliferation after exposure to senescence inducing agent. Using a high-throughput single-cell counting platform, we measured the proliferation of Hoechst 33342 stained cells over a three-day period at single-cell resolution (**Figure 1c**). Fibroblasts were treated with six different concentrations of bleomycin ranging from 0 (vehicle control) to 50 µM. On 4 GPa plastic substrates, cells followed a dose-dependent proliferation response (**Figure 1d**). At high doses (5 µM and 50 µM), proliferation was fully inhibited within 24h and stayed inhibited over the 3-day time course (**Figure 1d**). In a vehicle control (bleomycin-free) condition, cell proliferation increased >3.5-fold over the 3-day period for fibroblasts seeded on a 4 GPa microenvironment (**Figure 1d**, black curve).

Tissue culture plastic or glass substrates do not recapitulate the ECM stiffness of soft tissues. Therefore, we asked whether matrix stiffness changed the development of the senescence phenotype. In control conditions, fibroblasts seeded on type-I collagen-coated 0.5 kPa and 25 kPa PAA substrates showed an expected morphology (**Supplemental Figure 1a**) and roughly a 3-fold increase in cell number over the three-day period (black curves in **Figure 1e, 1f** corresponding to vehicle control) ^24–27^. However, when subjected to a senescence-inducing agent, bleomycin, primary fibroblasts showed significant changes in their proliferative response (**Figure 1e, f**). We observed a significant increase in the propensity for cells to enter a non-proliferative state on soft 0.5 kPa and stiff 25 kPa substrates relative to 4 GPa plastic. We also noted that complete inhibition of proliferation was observed at roughly 1/10^th^ the concentration required to arrest cell proliferation on plastic. On the softest matrix (0.5 kPa), fibroblasts treated with 250 nM bleomycin experienced a modest 44% increase in cell number over three days (**Figure 1f**) compared to nearly 200% for plastic (**Figure 1d**), a near total growth arrest at 1/20^th^ the required dose of plastic. These results were also supported in WI-38 embryonic lung fibroblast cell line cultured on various substrates and exposed to bleomycin (**Supplemental Figure 1d-e**).

Taken together, this data indicates that fibroblasts exposed to senescence-inducing drug bleomycin have a higher propensity towards entering a non-proliferative, senescent like state on softer substrates (0.5-25 kPa) compared to stiff tissue-culture plastic (**Figure, 1d-f, Supplemental Figure 1d-f**).

### Substrate stiffness regulates the rate of senescence induction

Since decreasing substrate stiffness promoted a senescence-associated proliferation arrest post induction (**Figure 1**), we asked whether substrate stiffness also changed the rate of senescence induction. Considering the proliferation experiments, we hypothesized that softer microenvironments would also be more conducive to senescence with an increased rate of senescence induction when exposed to bleomycin. To quantify this relationship between substrate stiffness and rate of senescence, we developed a mathematical model based on a simplified cell differentiation balance (**Figure 2a**). Our model operates under three fundamental assumptions 1) there are only two unique cell populations, *i.e.,* proliferating (non-senescent) and senescent; 2) cell death and pre-existing senescent cell populations are negligible, and 3) cell transitions from proliferative to senescence is irreversible. The growth constant *k_p_* for proliferating cells is a function of matrix stiffness. The growth constant *k_s_*for senescent cells is a function of both matrix stiffness and bleomycin concentration. Therefore, the total time-dependent number of cells *N*(*t*) can be represented by the equation (see derivation in

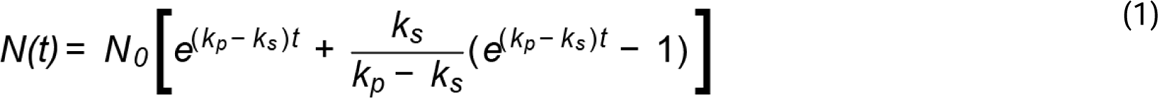

**Figure 2.**
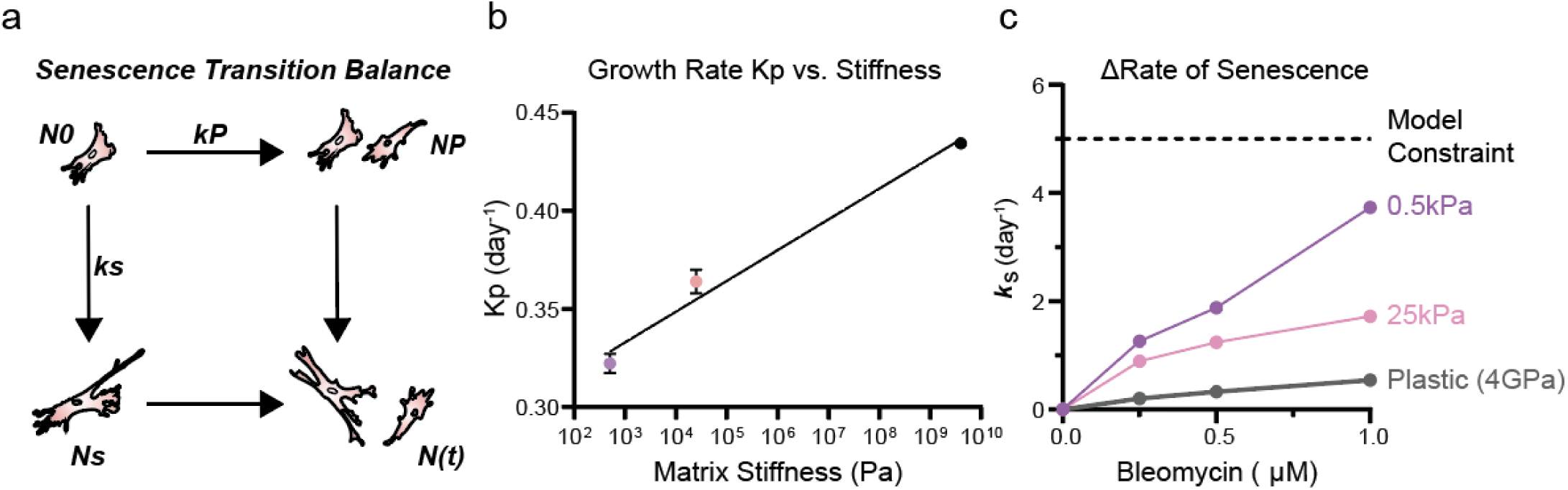
Substrate stiffness regulates the rate of senescence induction. (**a**) Balance illustrating the two potential outcomes for a fibroblast subjected to a senescence inducing microenvironment. (**b**) Numerically solving for experimental growth dynamics of unconstrained, drug free growth as a function of matrix stiffness shows higher rate of proliferation on stiffer substrates. (**c**) Rate of cellular senescence k_s_ on Plastic (4GPa), 25 kPa, and 0.5 kPa substrates as a function of bleomycin concentration, reveals the rate of onset for cellular senescence increases with decreasing substrate stiffness. Senescence transition model fit to 3 biological replicates each having 3-5 technical replicates.

Materials and Methods):

To determine the rate of proliferation *k*_p_ as a function of substrate stiffness, we used data generated from experimental growth dynamics, which showed a higher rate of proliferation on stiffer substrates (**Figure 2b**, **Supplemental Figure 2a-c**). To determine the rate of senescence induction *k_s_*, we numerically solved the equation for each substrate per bleomycin concentration (**Figure 1d-f, Supplemental Figure 2d**). Solving for *k_s_* revealed a stiffness-dependent rate of senescence induction (**Figure 2c**), with *k_s_* increasing as a function of decreasing substrate stiffness (highest *k_s_* on 0.5kPa and lowest on 4GPa) for all bleomycin concentrations tested. Taken together, these results suggest that cells on softer microenvironments transition to a senescence-like state at a higher rate relative to cells on stiffer microenvironments.

### Substrate stiffness modulates the expression of senescence biomarkers p16^INK4a^ and p21^WAF1/CIP1^

While proliferation arrest is a functional response to senescence induction, we asked whether other senescence defining features and biomarkers also changed with substrate stiffness. Senescence is typically associated with the overexpression of cell-cycle proteins p16^INK4a^ and p21^WAF1/CIP1^ and morphological changes that include increases in cell size and nuclear size^4,22,28^. To simultaneously quantify time-dependent changes in the expression of key senescence biomarkers and nuclear morphology at single-cell resolution, we employed an image-based high throughput cell phenotyping platform (htCP) (**Figure 3a-b**)^29,30^. As a proof of concept, we induced senescence in cells on 4GPa plastic substrates and investigated time-dependent changes in nuclear morphology and senescence biomarker expressions (p16^INK4a^, and p21^WAF1/CIP1^) for up to seven days post bleomycin induction (**Figure 3c**). We observed that the nuclear area of senescence-induced fibroblasts increased approximately 1.65-fold, from an average nuclear area of 251 µm^2^ on day 1 to 414 µm^2^ on day 7 (**Figure 3d**). Based on measurements of nuclear aspect ratio (long axis length divided by short axis length) and shape factor (SF=4πA/p^2^), we observed that cells adopted a more elongated and less circular nuclear morphology (**Supplemental Figure 3a-c**). In addition to the expected increase in nuclear size following senescence induction, we observed a characteristic increase in the expression of p21^WAF1/CIP1^ and p16^INK4a^. The total nuclear content of p21^WAF1/CIP1^ and p16^INK4a^ increased by ∼3-fold and ∼2-fold, respectively, when normalized to baseline non-senescent vehicle control (**Figure 3e-f**), thereby corroborating the senescent phenotype.

**Figure 3.**
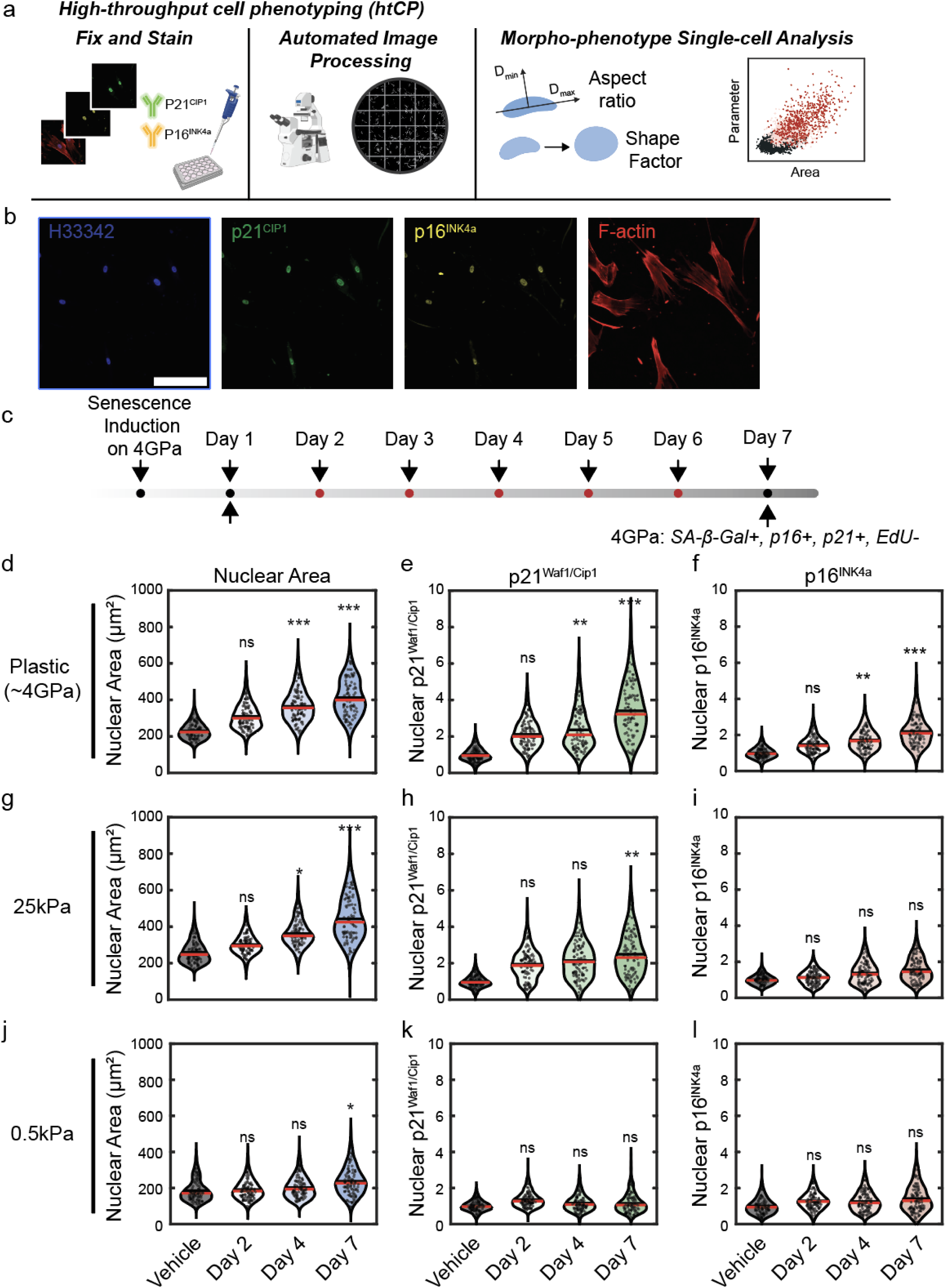
Substrate stiffness modulates the expression of senescence biomarkers p16^INK4a^ and p21^WAF1/CIP1^. (**a**) High throughput cell phenotyping (htCP) platform allows for identification of senescence associated morphological and molecular phenotypes at a single cell resolution. (**b**) Representative 10X immunofluorescent images of GT22 fibroblasts seven days after senescence induction stained for p16^INK4a^ and p21^WAF1/CIP1^.Scale bar 200µm (**c**) Induction workflow for quantification of senescence associated molecular biomarker kinetics. Senescent cells experiencing a 4GPa plastic substrate show increased (**d**) nuclear area (µm^2^), (**e**) nuclear expression of p21^WAF1/CIP1^ and (**f**) nuclear expression of p16^INK4a^ with time. Senescent cells on 25kPa substrates show increased (**g**) nuclear area but reduced (**h**) nuclear p21^WAF1/CIP1^ and (**i**) nuclear p16^INK4a^ with time. 0.5kPa fibroblasts show (**j**) decreased nuclear growth and negligible expression of (**k**) nuclear p21^WAF1/CIP1^ and (**l**) nuclear p16^INK4a^ with time. Single cell immunofluorescence data evaluated via one-way ANOVA, with Tukey multiple comparison test to vehicle control. Statistics evaluated based on mean of N=3 biological replicates (***P <0.001, **P < 0.01, *P < 0.05).

To investigate whether substrate stiffness changed the expression of the above senescence biomarkers, we seeded non-senescent fibroblasts on 4 GPa plastic substrates and induced senescence using bleomycin (see Methods). At 24 h post-induction, induced cells were dissociated and seeded onto collagen-I coated substrates of 4 GPa, 25 kPa and 0.5 kPa stiffness (**Figure 3c**). Bleomycin-treated cells on 25 kPa showed near identical nuclear morphological response as cells on 4 GPa (**Figure 3g**, **Supplemental Figure 3a-c**). However, we observed significant differences in the time-dependent changes in senescence biomarkers on softer substrates. On 25 kPa substrates, nuclear p21^WAF1/CIP1^ expression increased ∼2-fold relative to non-senescent baseline cells, which was less than the ∼3-fold change for cells on 4 GPa substrates (**Figure 3h**). Interestingly, we observed no significant change in nuclear p16^INK4a^ at day seven relative to baseline non-senescent cells (**Figure 3i**). This result was amplified for cells on 0.5 kPa substrates, with a moderate but significant increase in nuclear size at day seven relative to non-senescent cells (∼1.4-fold increase) (**Figure 3j**) and no significant changes in either p21^WAF1/CIP1^ (**Figure 3k, Supplemental Figure 3d**) or p16^INK4a^ (**Figure 3l, Supplemental Figure 3e**) at day seven. This lack of a conventional senescence phenotype based on protein biomarkers was similarly observed in WI-38 embryonic lung fibroblasts (**Supplemental Figure 3g-o**) and supported by SA-β-gal, with senescent cells on 4 GPa substrates showing strong SA-β-Gal staining and reduced positivity for cells on 25 kPa and even less for cells on 0.5 kPa substrates (**Supplemental Figure 1c**). Hence, conventional senescence protein-based biomarkers do not appear in senescent cells on soft substrates even though these cells have undergone complete proliferation arrest (**Figure 1**). Taken together, these results suggest that the senescence phenotype measured via biomarkers is mechanosensitive.

### Focal adhesion signaling regulates mechanoresponsive p21^WAF1/CIP1^ expression in senescent fibroblasts

A mechanosensitive response to senescence induction presents significant challenges in identifying senescence in tissues, as tissue stiffness varies widely across tissue-types, and during aging and disease^31^. To further investigate the lack of p16^INK4a^ and p21^CIP/WAF1^ expression for senescence-induced cells on soft 0.5 kPa substrates, we asked whether the observed mechanoresponsive phenotypes were reversible. To assess reversibility, we took both baseline non-senescent cells and bleomycin-induced cells from their 0.5 kPa and 4 GPa substrates where they had been cultured for 6 days and reseeded them onto 4 GPa plastic substrates (**Figure 4a**). Results showed that 48 h after transfer (*i.e.,* nine days after initial induction) to 4 GPa substrates, senescent cells cultured on 0.5 kPa and 4 GPa substrates both recovered to exhibit characteristically larger nuclear morphologies (**Figure 4b**) and high expression of p16^INK4a^ and p21^CIP/WAF1^ (**Figure 4c, d**).

**Figure 4.**
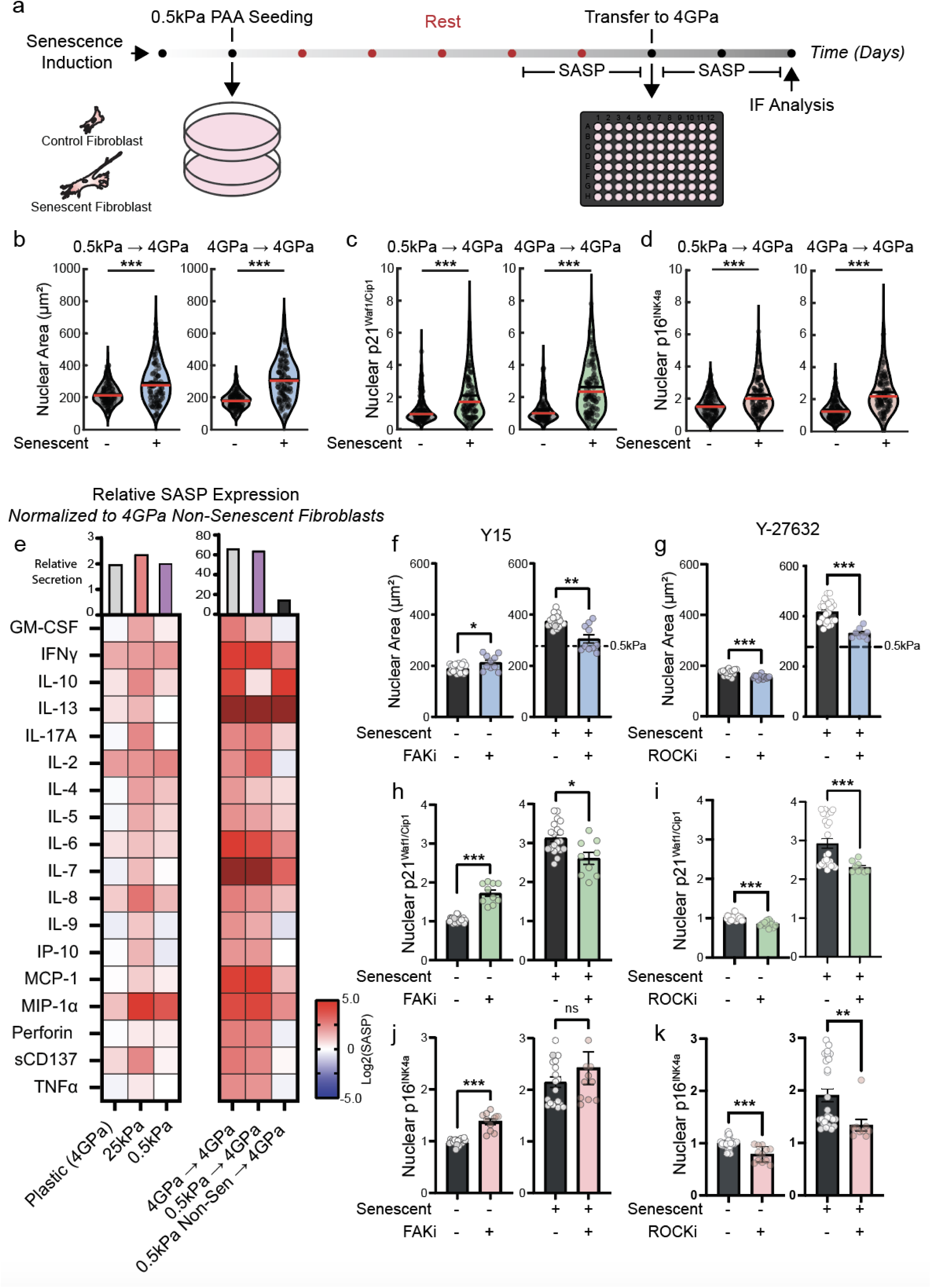
Focal adhesion signaling regulates mechanoresponsive p21^WAF1/CIP1^ expression in senescent fibroblasts. (**a**) Transfer of senescent fibroblasts to a non-physiological plastic substrate revealed a (**b**) recovery of the senescent morphological and (**c**) p21^WAF1/CIP1^ (**d**) p16^INK4a^ phenotype. (**e**) Multiplex cytokine panel revealed SASP expression of senescent cells experiencing 0.5kPa, 25kPa, and 4GPa substrates show unique, but upregulated SASP profiles. When transferred from soft 0.5 kPa subtrates to a 4GPa (plastic) substrate, 0.5kPa cells match the SASP of 4GPa (plastic). Normalized to non-senescent 4GPa condition before and after transfer. Deep red indicated >5x log2(fold change) in relative expression. Crown illustrates change in total protein secretion relative to 4GPa non-senescent condition. (**f**) Small molecule inhibition of the focal adhesion pathway via Y-15 (FAKi, 10µM) decreased nuclear area in senescent fibroblasts on plastic 4GPa to comparable 0.5kPa conditions. (**g**) inhibition of the focal adhesion pathway via Rho-Associated protein kinase (ROCK) inhibitor Y-27632 (ROCKi, 10µM) decreased in nuclear area in senescent fibroblasts on plastic substrates back to that of senescent fibroblasts on 0.5 kPa substrates. (**h**) Treatment with FAKi and (**i**) ROCKi decreased senescent associated p21 expression in senescent fibroblasts. (**j**) FAKi increased nuclear p16^INK4a^ in non-senescent fibroblasts but had no response on senescent conditions (**k**) ROCKi had no effect on p16^INK4a^ expression in non-senescent fibroblasts but decreased significantly in senescent fibroblasts. Error bars represent SEM of biological variation, statistics evaluated via Welch’s t test, 3 biological replicates each having 3-5 technical replicates. (***P <0.001, **P < 0.01, *P < 0.05).

As observed with p16^INK4a^, and p21^WAF1/CIP1^ expressions and SA-β-Gal staining, the cytokine secretion profile of senescent cells also depended on substrate stiffness (**Figure 4e, left panel**). Forty-eight hours post transfer, senescent cells on that had been cultured on 0.5kPa substrates had a near identical inflammatory response in cytokines like IL-6, IL-8, and IFNγ compared to 4GPa (**Figure 4e, right panel**). This highly inflamed SASP response was not reflected to the same magnitude in non-senescent 0.5kPA conditions (**Figure 4e, right panel**). Together, the recovery of a conventional senescent phenotype following transfer to a 4 GPa substrate indicates that the expression of senescence biomarkers is mechanosensitive.

To further confirm the mechanosensitive nature of the senescence phenotype, we treated cells with inhibitors of modulating cytoskeletal tension^22,32^. Focal adhesions (FAs) are organized multi-protein structures that transmit mechanical forces from the ECM/substrate to the cytoskeleton. FAs regulate a multitude of processes including cell morphology, cell movements, and proliferation^33,34^ and are composed of several subunits, including focal adhesion kinase (FAK)^35^. To test whether cells treated with a FAK inhibitor exhibited a similar phenotype to cells seeded on soft 0.5 kPa substrates, we treated senescent and non-senescent cells on 4 GPa substrates with FAK inhibitor Y15 (**Supplemental Figure 4a**). We observed that in senescent cells treated with 10 μM Y15, the nuclear area decreased to a size comparable to cells seeded on 0.5 kPa substrates (**Figure 4f**). This loss of nuclear spreading was accompanied by a decrease in nuclear expression of p21^WAF1/CIP1^ (**Figure 4h**), although p16^INK4a^ remained unchanged (**Figure 4j**). Both p21^WAF1/CIP1^ (**Figure 4h**) and p16^INK4a^ (**Figure 4j**) increased in non-senescent cells treated with FAKi, a result indicative of a slowed cell-cycle consistent with FAK inhibition^36^. Collectively these results suggest that cytoskeletal tension and mechanotransduction may be responsible for the lack of biomarker phenotypes observed on softer substrates.

FAK signaling is associated with acto-myosin contractility via the Rho-kinase (ROCK), which regulates the phosphorylation of myosin light chain ^37–39^. To test whether inhibiting ROCK yielded a similar result as FAK inhibition, we treated both senescent and non-senescent cells with ROCK inhibitor 10 μM Y-27632 (**Supplemental Figure 4b**). We observed that senescent cells seeded on 4 GPa substrates treated with ROCKi exhibited a nuclear area comparable to cells seeded on 0.5 kPa substrates (**Figure 4g**), and a significant decrease in nuclear expression of both p21^WAF1/CIP1^ (**Figure 4i**) and p16^INK4a^ (**Figure 4k**), with a less than 5% change in the expression of p21^WAF1/CIP1^, and p16^INK4a^ for non-senescent cells treated with Y-27632 (**Figure 4g, I, k**).

Collectively, these results indicate that senescent cells are mechanosensitive and that external mechanical cues can regulate nuclear morphology and senescence biomarker expression via outside-in mechanical signaling involving focal adhesions and acto-myosin contractility.

### Mechanosensitive regulation of CDKIs p16^INK4a^ and p21^WAF1/CIP1^ is initiated at the RNA level

Given that senescent cells on soft substrates do not show the characteristic increases in biomarker expressions at the protein level (**Figure 3**), we wanted to understand where the senescence phenotype was being regulated. Molecular expression of p21^WAF1/CIP1^ and p16^INK4a^ are tightly controlled by competing rates of production and degradation^40,41^. Hence, we asked whether the mechanosensitive character of the senescence phenotype occurred at the transcription, translation, or post-translational levels. Using qPCR, we measured how the gene expression of p21 (*CDKN1a)*, p16 (*CDKN2a)* and core SASP gene IL-6 (*IL6)* changed with substrate stiffness. *CDKN1a* and *CDKN2a* were both upregulated in senescent fibroblasts on 4 GPa substrates (**Figure 5a-c**). On 25 kPa substrates, *CDKN1a, CDKN2A* and *IL6* were all upregulated in senescent cells, although to a slightly lesser extent relative to cells on 4 GPa substrates (**Figure 5a-c**). For cells on 0.5 kPa substrates*, CDKN1a* and *IL6* expressions were higher for senescent cells, however, the expression of *CDKN2a* was unchanged relative to non-senescent cells (**Figure 5b**). These results suggest that while bleomycin-treated cells on soft substrates activate the senescence program at the RNA level, however, the cellular response to this soft substrate suppresses senescence biomarkers at the translation or post-translational level.

**Figure 5.**
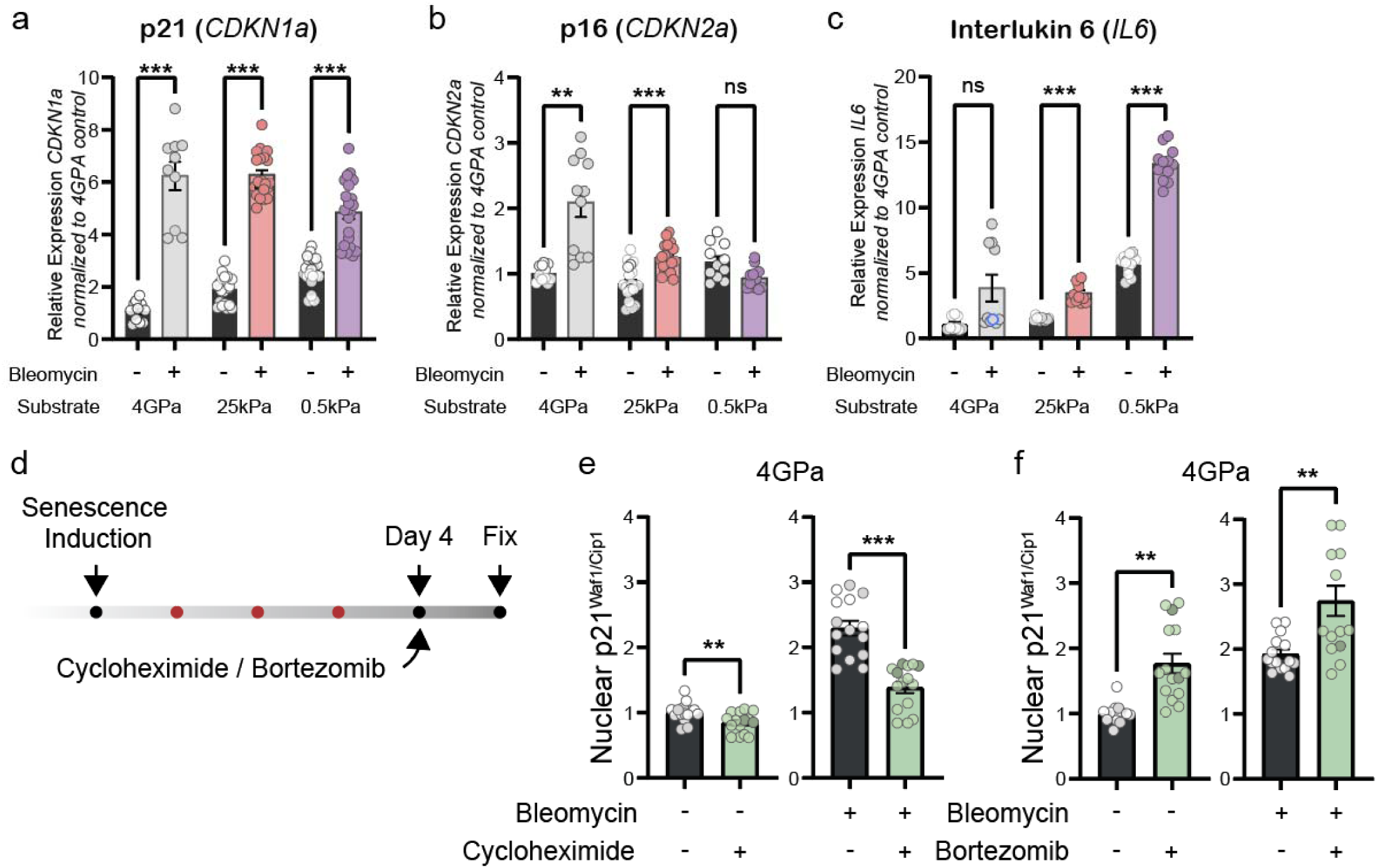
Mechanosensitive regulation of CDKIs p16^INK4a^ and p21^WAF1/CIP1^ is driven at the translational level. (**a**) *CDKN1a* gene expression decreases in senescent fibroblasts cultured on 25 kPa, and 0.5 kPa conditions relative to plastic control. (**b**) *CDKN2a* gene expression between senescent and non-senescent fibroblasts converge at soft substrates. (**c**) IL6, a core SASP gene is upregulated on soft substrates and increased in senescent conditions. (**d**) Workflow for small molecule inhibition of protein synthesis (Cycloheximide) and protein degradation (Bortezomib) of early senescent fibroblasts. (**e**) Early senescent fibroblasts treated with 1µM Cycloheximide reverted the p21^CIP/WAF1^ protein phenotype to that of vehicle control. (**f**) treatment with 100nM Bortezomib increased p21^WAF1/CIP1^ in both non-senescent and senescent cell populations. Error bars represent SEM of biological variation, statistics evaluated via Welch ANOVA with Dunnett correction for multiple comparisons, 3 biological replicates each having 3-5 technical replicates. (***P <0.001, **P < 0.01, *P < 0.05).

To test how translation and post-translation modifications altered the senescence phenotype, we treated senescence-induced cells four days after bleomycin induction on 4 GPa (with elevating expression of p16^INK4a^ and p21^WAF1/CIP1^) with 1µM cycloheximide to inhibit protein translation^42^ (**Figure 5d, Supplemental Figure 4c**). Inhibition with cycloheximide during the onset of the senescence phenotype decreased nuclear p21^WAF1/CIP1^ to levels observed in non-senescent conditions (**Figure 5e**) and was accompanied by a decrease in nuclear area (**Supplemental Figure 4d**) with no change in p16^INK4a^ expression (**Supplemental Figure 4e**). These results indicate that inhibiting protein translation during the development of the senescence phenotype leads to a decrease in p21^WAF1/CIP1^ and nuclear area and a lack of conventional senescence biomarker expression, as observed in cells on soft 0.5 kPa substrates. Blocking protein degradation using 100nM Bortezomib showed no change between senescence and non-senescent fibroblasts both in morphology, and molecular expression of p21^WAF1/CIP1^ and p16^INK4a^ (**Figure 5f, Supplemental Figure 4f-h**. Together, this data suggests that the decreased p21^WAF1/CIP1^ protein expression in cells on soft substrates can be attributed to reduced protein translation and little change in degradation. While p21^WAF1/CIP1^ shows a significant response, p16^INK4a^ on the other hand did not show significant change.

## Discussion

Cellular senescence is an established driver of aging and is associated with a range of pathologies, including fibrosis, cancers, diabetes, and Parkinson’s^1,41^. Here we demonstrate the conventional senescent phenotype based on molecular biomarkers is mechanosensitive. Using a dermal fibroblast model of senescence, we show that substrate stiffness regulates both cellular proliferation arrest and molecular biomarker expression in the presence of a senescence-inducing drug. Modeling of proliferation dynamics reveals that the rate of senescence is stiffness-dependent and highest on soft substrates. The expressions of cyclin-dependent kinases inhibitors - p21^WAF1/CIP1^ (*CDKN1a*) and p16^INK4a^ (*CDKN2a*), which are commonly used as biomarkers of senescence, are mechanosensitive. These CDKIs are activated transcriptionally during the senescence program but regulated at translation through myosin contractility and focal adhesion signaling.

This work has potential implications for the identification of senescent cells in tissue. A direct measurement of cell proliferation which can be done at single-cell resolution *in vitro*, cannot be readily done *in vivo*. Hence, assessments of senescence in vivo rely solely on proxies of senescence, *i.e.,* senescence biomarkers, mainly protein based. Given our observations, local changes in tissue stiffness may both drive the onset of senescence and the ability to measure the senescence phenotype. While it is widely accepted that there is no single biomarker for senescence, intracellular expression of p16^INK4a^ and p21^WAF1/CIP1^ are currently the *de facto* method for identifying senescence in tissue sections^4,7^. In the absence of better molecular signatures, careful interpretation of these protein-based biomarker expressions is needed, particularly in mechanically soft tissues such as the brain and pancreas^8^. Our work suggests that at the protein level senescence phenotypes do not manifest on softer matrices yet at the RNA level, the process is transcriptionally initiated. In addition, recent work has shown that γH2AX and SASP expression also decrease with stiffness, even while maintaining transcriptional activation^43,44^. The fact that transcriptional programs of senescence are conserved bodes well for RNA-based signatures to identify senescence in softer substrates/environments. As such, approaches using resources like SenMayo and SenSIG, which are RNA-based signature may be useful^45,46^.

Interestingly, in addition to paracrine signaling through SASP, senescence is a mechanically active phenotype known to increase tissue stiffness in response to wound healing^42^. This biophysical response suggests that senescent cell-ECM interactions may ultimately accentuate biomarker expression. Spatially resolved mechanical information obtained in combination with current proteome and transcriptome strategies might elucidate why senescence biomarker staining yields such heterogenous results. Unfortunately, mechanical mapping of precious tissues presents many logistical and technical hurdles, presenting a current limitation. However, given the various pathological pathways associated with substrate stiffness, future work is required to resolve the detailed mechanisms behind how senescent cells sense and respond to mechanical stimuli, and how to effectively identify these senescent cells in mechanically soft microenvironments.

## Data Sharing

All data critical to the findings are presented within the manuscript and the supplementary documents.

## Acknowledgements.

This work was supported through grants from the National Cancer Institute (U54CA143868 and U54CA268083 to PW, DW), National Cancer Institute (UG3CA275681 to PW, DW, JMP), the National Institute of Arthritis and Musculoskeletal and Skin Diseases (U54AR081774 to DW, PW, JMP) and the National Institute on Aging (U01AG060903 to DW, JMP), The Johns Hopkins University Older Americans Independence Center of the National Institute on Aging under award number P30AG021334 (JMP), Start-up funds from the Department of Biomedical Engineering and the Whiting School of Engineering at Johns Hopkins University (JMP).

## Contributions

BS, DW, JMP conceived, developed, and designed the study. BS, FY, DT, JB, HK, JE, PK, NM performed experiments. SS, DW conceived and developed the senescence model. BS, PRN, PW, LG, JW, DW, JMP developed model systems and provided resources. BS, FY, KH performed formal analysis. BS, PRN, DW, JMP wrote and edited the manuscript with input from all co-authors. JMP, DW supervised study and secured funding.

## MATERIALS AND METHODS

### Cell Line Culture

Primary dermal fibroblast cell lines GT22 were obtained from The Baltimore Longitudinal Study of Aging (BLSA). Fibroblast cells were cultured in DMEM (Corning, 10-013-CV) supplemented with 15% (v/v) fetal bovine serum (FBS, Corning, 35-010-CV) and 1% (v/v) penicillin streptomycin (P/S, Millipore Sigma, PO781). Primary Fibroblasts were maintained and used at or below six passages from original cell line transfer. Immortalized, embryonic normal lung fibroblast cell line WI-38 (ATCC, CCL-75) was cultured in high-glucose DMEM (Gibco, 10-566-016) supplemented with 10%(v/v) FBS and 1% (v/v) P/S. Cell lines were maintained at 37°C and 5% CO_2._

### Polyacrylamide Gel Casting

6-well and 96-well Collagen-I coated polyacrylamide (PAA) gels of 0.5 kPa and 25 kPa elastic moduli were purchased commercially from Matrigen LLC. 24-well PAA gels of 0.5 kPa, 25 kPa and were cast in-house as described in^47^. Briefly, 40% (w/v) Acrylamide (Sigma-Aldrich, A3553) was mixed with 2% (w/v) N-N’-Methylenebisacrylamide (Sigma-Aldrich, M7279), DPBS for a final concentration of 10% acrylamide. Bis-acrylamide was varied to control elastic modulus. 10% Ammonium Peroxydisulfate (1:100) (Bio-Rad, #1610700) and TEMED (1:1000) (Bio-Rad, #1610800) were mixed with acrylamide-bis-Acrylamide solution. The solution was mixed, transferred, and plated between two siliconized (Sigma-Aldrich, SL2) glass casting plates. The gels polymerized for 45 minutes, at which point desired gel sized were cut, washed, and moved to 24-well plates. PAA hydrogels were functionalized with 0.2mg/mL Sulfo-SANPAH (ProteoChem, C1111) for 10 minutes under 365nm UV lamp (36W, 7cm) and coated with 0.05 mg/mL Type I Collagen (Corning, 354236) overnight at 2-8°C. Commercial and casted gels were gently washed 3X with DPBS immediately before use.

### Senescence Induction

The senescent phenotype was induced as described previously ^22^. Briefly, ∼1×10^5^ primary dermal fibroblasts were seeded in 100mm dishes and allowed to adhere overnight. Fibroblasts were then exposed to 50µM Bleomycin (Selleckchem, S1214) resuspended in a 1X DPBS (Corning, 21-031-CV) vehicle for 4hr. Post senescence induction, cells were washed 3X with DPBS and allowed to recover overnight from initial point of exposure. Post induction, fibroblasts were trypsinized, counted, and re-seeded on substrates of interest.

### Cell Proliferation

Cell proliferation was measured using an in-house high throughput-cell counting platform. In brief, fibroblasts were seeded at a density of 1000 cells/well in a 96 well black-walled plates containing an ECM functionalized polyacrylamide gel of varying elastic modulus – 0.5 kPa, 25 kPa or collagen coated 4GPa (glass). Primary fibroblasts were treated with 6 different titrations of bleomycin (Figure 1). At the end of induction (4hr), an initial cell count cell was conducted using cell-permeable DNA stain Hoechst 33342 (Invitrogen, H3570, 1:1000, 10µg/mL). Whole wells were stained and imaged in triplicate every day for three days using a 2X objective (Nikon, MRL00022) on a Nikon TI-Eclipse equipped with a Hamamatsu ORCA-Flash 4.0 CMOS camera and Lumencor SPECTRA X LED light source. WI-38 Fibroblasts were stained with live-cell fluorophore SPY555-DNA (Cytoskeleton, CY-SC201, 1:500), and imaged daily for three days following the same imaging protocol. A single-cell count was obtained via a local maxima bright spot detection and filtered using percentile filtering to remove extraneous noise such as debris, PAA gel artifacts, collagen in MATLAB.

### Modeling the rates of senescence induction

To generate the senescence transition curves, we derived a model that aimed to capture the rate of onset for cellular senescence in the presence of a senescence inducing drug (Bleomycin). Our model operates under three assumptions 1) there are only two unique cell populations – proliferating and senescent; 2) cell death and preexisting senescent cell populations are negligible, and 3) transition towards senescence is irreversible. The growth constant *k_p_* for proliferating cells is a function of only matrix stiffness. While the growth constant *k_s_* for senescent cells is a function of matrix stiffness and drug concentration. We note that proliferation and metabolism are intertwined when using analogs of proliferation (MTT and PrestoBlue), to remove dependence on metabolic activity, we directly counted cell nuclei ^48^. Therefore, the mean field equations can be written as:

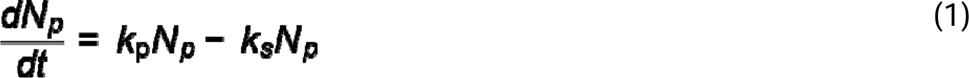

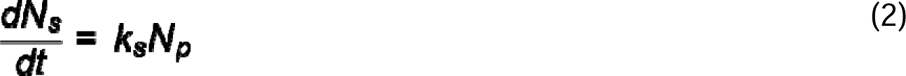

Integrating each equation yields the proportion of proliferating and senescent cells (*Ns,t*=0=0, *Np,t*=0=*N*0)

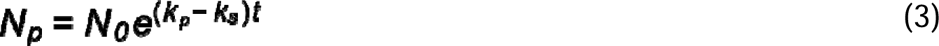

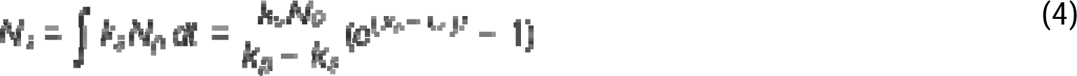

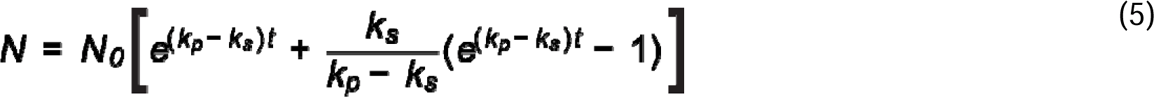

Where the total number of cells can be represented by:

Vehicle growth kinetics data was LN transformed and fit with a linear regression to extract a drug free growth constant (*k_p_*) for each substrate of interest. Cell proliferation data for all substrates and drug conditions were fit using a least squares regression the senescence transition model solving for *k_s_* at each drug concentration. Model constraints include the following: 1) *k_p_* were assumed constant and constrained to a constant for each substrate. 2) Independent variable *k*_s_ varies with drug concentration and matrix stiffness but promptly reaches total proliferation arrest at high drug concentrations (>1µM on soft substrates). Therefore, *k_s_*was constrained between 0 (no drug) and 5 (total proliferation arrest). The upper bounded constraint of *k_s_* represents a slope within 10% of normality.

### Immunofluorescence

Immunofluorescence was carried out as described previously. Briefly, 24hr post induction, Fibroblasts were seeded at a density of 400 cells/well in a 96 well black-walled plates containing an ECM functionalized polyacrylamide gel of varying elastic modulus – 0.5 kPa, 25 kPa, and 4 GPa. For kinetics experiments, cells were fixed every day for seven days with 4% (w/v) Paraformaldehyde (PFA) (Sigma-Aldrich, 158127) DPBS solution. Fixed cells were washed 3X in DPBS. At the end of seven-day period, cells were permeabilized for 10 minutes using a 1% (v/v) Triton X-100 (Sigma-Aldrich, X100) DPBS solution, and washed 2X with DPBS. Permeabilized cells were stored in a 0.1% (w/v) Sodium Azide (Sigma-Aldrich, SX0299) in DPBS solution.

For antibody staining, permeabilized cells were blocked for 30 minutes at room temperature (RT) in 5% (v/v) normal goat serum (NGS) (Cell Signaling Technology, #5425) DPBS solution. Primary antibodies were incubated separately to avoid potential aggregation. Cells were first incubated overnight at 4°C and 300rpm with p21^WAF1/CIP1^ (Santa Cruz, SC-817, 1:200, 1.0µg/mL), suspended in a 2.5% NGS solution. The following day, cells were washed 3X with DPBS, and incubated with p16^INK4a^ (ABCAM, AB108349, 1:250, 5.80µg/mL) suspended in 2.5% NGS solution for 4h at RT at 300rpm on an orbital shaker. Cells were washed 3X with DPBS, and counterstained with Alexa-Fluor conjugated secondary antibodies (Invitrogen A11001 & A11011, 1:500, 4µg/mL) for 2 h at RT. Cells were washed 3X with DPBS and stained with small molecule DNA stain Hoechst 33342 (Invitrogen, H3570, 1:1000, 10µg/mL), and Phalloidin 647 Plus (Invitrogen, A30107, 1:400) to visualize nuclear and F-actin. Stained cells were imaged at 10x on Nikon Ti-Eclipse equipped with a Hamamatsu ORCA-Flash 4.0 CMOS camera.

### High throughput cell phenotyping (htCP) immunofluorescence quantification

Immunofluorescent images were quantified using a high throughput cell phenotyping (htCP) platform^29^. htCP outputs both single cell morphological and molecular information for each cell in a stitched frame field of view. Single-cell data was further refined based on a rigorous gating strategy of nuclei to filter outlier populations such as – optical aberrations, mis-segmentations, debris, antibody aggregates, and optical edge effects (halos). Gating results were matched via overlay of identified nuclei on original fluorescent images. Protein biomarker integrated intensity data was normalized based on the mean of each vehicle control condition, grouped with other biological replicates, and plotted in MATLAB.

### Recovery Experiment

For phenotype recovery experiments, GT22 fibroblasts were plated on 100mm dishes and allowed to adhere overnight. Cells were induced with 50µM bleomycin or vehicle control. 24hr later, cells were typsinized, counted, and seeded on 0.5 kPa 100mm dishes at a density of 3×10^5^ cells/dish. Cells were allowed to rest on the 0.5 kPa PAA substrate for 6 days at which point plates were washed, cells dissociated via 10X TrypLE and seeded on collagen coated 96-well 4GPa plates at a density of 500 cells/well. Fibroblasts were allowed to recover for 24 or 72hr – at which point cells were fixed, permeabilized, and stained with p21^WAF1/CIP1^ (Santa Cruz, SC-817, 1:200, 1.0µg/mL), p16^INK4a^ (ABCAM, AB108349, 1:250, 5.80µg/mL), Hoechst 33342 (Invitrogen, H3570, 1:1000, 10µg/mL), and Phalloidin 647 Plus (Invitrogen, A30107, 1:400). Plates were imaged at 10x (MRF00101) on Nikon Ti-Eclipse equipped with a Hamamatsu ORCA-Flash 4.0 CMOS camera. Molecular expression data was quantified via htCP.

### Focal Adhesion kinase (FAK) and Rho-associated protein kinase (ROCK) Inhibition

For mechanosensing experiments, GT22 fibroblasts were plated on 100mm dishes and allowed to adhere overnight. Cells were induced with 50µM bleomycin or vehicle control for 4 h, washed and the senescent phenotype allowed to develop for seven days. Seven days later, cells were typsinized, counted, and seeded at a density of 400 cells/well on collagen coated plastic plates and allowed to adhere overnight. Vehicle control, and Senescent induced cells were treated for 48hr with a series of titrations of Y15, and Y-27632 (50µM, 20µM, 1µM, 500nM, 100nM). After 48hr, cells were fixed, permeabilized, and stained with p21^WAF1/CIP1^ (Santa Cruz, SC-817, 1:200, 1.0µg/mL), p16^INK4a^ (ABCAM, AB108349, 1:250, 5.80µg/mL), Hoechst 33342 (Invitrogen, H3570, 1:1000, 10µg/mL), and Phalloidin 647 Plus (Invitrogen, A30107, 1:400). Plates were imaged at 10x (MRF00101) on Nikon Ti-Eclipse equipped with a Hamamatsu ORCA-Flash 4.0 CMOS camera. Molecular expression data was quantified via htCP workflow.

### Proteosome and Translation Inhibition

For Cycloheximide and Bortezomib treatments, GT22 fibroblasts were plated on 100mm dishes and induced with 50µM bleomycin or vehicle control. 24hr later, cells were typsinized, counted, and seeded at a density of 400 cells/well on collagen coated black walled plates and allowed to adhere overnight. Vehicle control, and Senescent induced cells were treated for 24hr with a series of titrations of Cycloheximide (50uM, 20uM, 1uM, 500nM) or Bortezomib (100nM, 10nM, 1nM, 500pM). 24hr after Cycloheximide/Bortezomib addition cells were fixed, permeabilized, and stained with p21^WAF1/CIP1^ (Santa Cruz, SC-817, 1:200, 1.0µg/mL), p16^INK4a^ (ABCAM, AB108349, 1:250, 5.80µg/mL), Hoechst 33342 (Invitrogen, H3570, 1:1000, 10µg/mL), and Phalloidin 647 Plus (Invitrogen, A30107, 1:400). Plates were imaged at 10x (MRF00101) on Nikon Ti-Eclipse equipped with a Hamamatsu ORCA-Flash 4.0 CMOS camera. Molecular expression data was quantified via htCP workflow.

### SA-B-Gal Staining

GT22 fibroblasts were plated on 100mm dishes and induced with 50uM bleomycin or vehicle control. 24hr later, cells were trypsinized, counted, and seeded at a density of ∼1500 cells/well on 24 well glass bottom plates containing: collagen coated glass, 0.5 kPa and 25 kPa PAA hydrogels. Fibroblasts were allowed to recover for seven days, at which point cells were fixed and stained for SA-B-Galactosidase (Millipore Sigma, CS0030). Plates were incubated at 37°C without CO_2_ for 48h. Brightfield images were taken at 10x (MRF00101) on Nikon Ti-Eclipse equipped with a Hamamatsu ORCA-Flash 4.0 CMOS camera.

### EDU Staining

Bleomycin and vehicle induced cells were seeded at a density of: 6×10^3^/3×10^3^ (Bleo/Vehicle) cells/well on 0.5 kPa PAA gels and 3×10^3^/1.5×10^3^ (Bleo/Vehicle) cells/well on 25 kPa PAA hydrogels and collagen coated glass plates. Cells were allowed to recover for six days, at which point fibroblasts were treated for 24 h with 10µM EdU solution (Invitrogen, C10337). 24 h after EdU treatment, cells were fixed with 4% PFA 10 minutes, washed 3X with 3% BSA in PBS, permeabilized with 0.5% Triton X-100 in DPBS for 20 minutes and stained for 30 minutes each with Click-iT reaction cocktail containing Alexa Fluor 488 Azide and 5ug/ml Hoechst 33342 (Invitrogen, H3570). The plate was imaged at 10x (MRF00101) on Nikon Ti-Eclipse equipped with a Hamamatsu ORCA-Flash 4.0 CMOS camera.

### RT-PCR

Total RNA from fibroblasts subjected to different stiffness conditions and drug treatments was isolated and purified using the RNeasy Plus Mini Kit (QIAGEN). cDNA was synthesized from 1 µg RNA with the iScript cDNA Synthesis Kit (Bio-Rad). Real-time PCR (RT-PCR) reactions were performed in a 384 well plate using iTaq Universal SYBR Green Supermix (Bio-Rad) and a thermal cycler (CFX384^TM^ Real-Time System, Bio-Rad). Normalization for each condition was conducted using the geometric mean of the RPS18, RPL13A, and H3F3A housekeeping Cq values, and relative expression was determined using the △△Ct method. The list of primers used in the study are below.

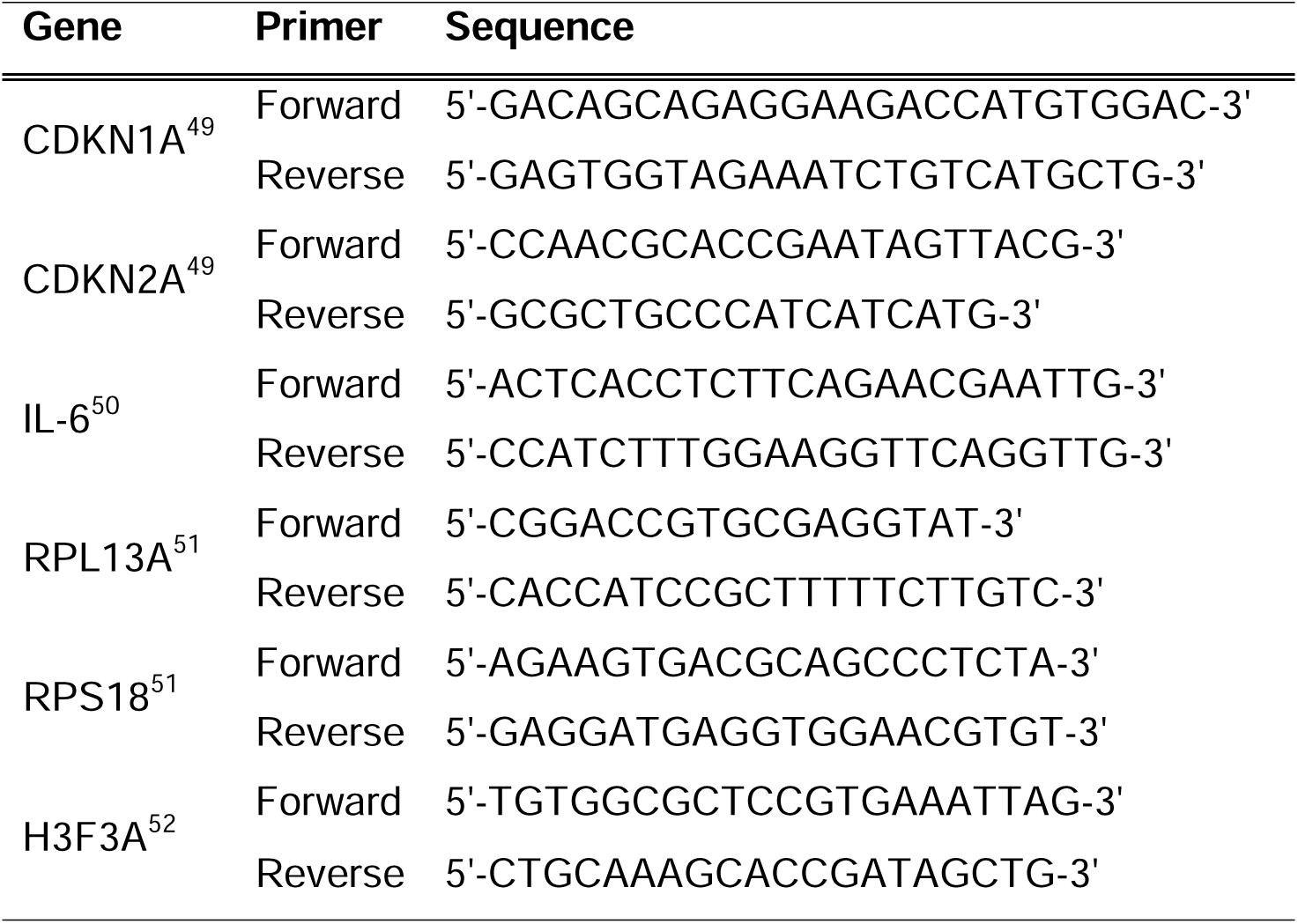

### Secretomic analysis of senescent fibroblasts before and after cell transfer

Senescent and non-senescent fibroblasts induced on plastic (4 GPa) substrates were seeded on 4GPa plastic, 25 kPa, and 0.5 kPa substrates 24 h after senescence induction at a density of 100,000 cells (plastic 4 GPa), and 200,000 cells (25 kPa, and 0.5 kPa) in 10mL medium per 10cm dish. For bulk cytokine analysis, medium was refreshed 5 days after senescence induction, condition medium was collected 48 h later (7-days). At which point cells were dissociated using 10X TypLE, counted, and transferred to corresponding substrate of interest. Condition medium was once again collected 48hr later. Conditioned medium was run on an IsoSpark (Bruker) using a preconfigured analyte panel (Codeplex, Human Adaptive Immune). Senescent and non-senescent condition medium was run in duplicate following the manufacturer protocol. Data was background corrected to a medium blank and processed using proprietary ISOSPEAK software. Senescent conditions were normalized to plastic control conditions before and after transfer. Heatmaps were generated in MATLAB.

### Statistical analysis

Mean values +/− SEM were calculated and plotted using GraphPad PRISM or MATLAB. Statistical tests used are denoted in figure captions. Significance is indicated in graphs using Michelin grade scale (***P <0.001, **P < 0.01, *P < 0.05).

