## Supplemental Figure for "Substrate stiffness modulates the emergence and magnitude of senescence phenotypes in dermal fibroblasts"

Supplemental Figure 1

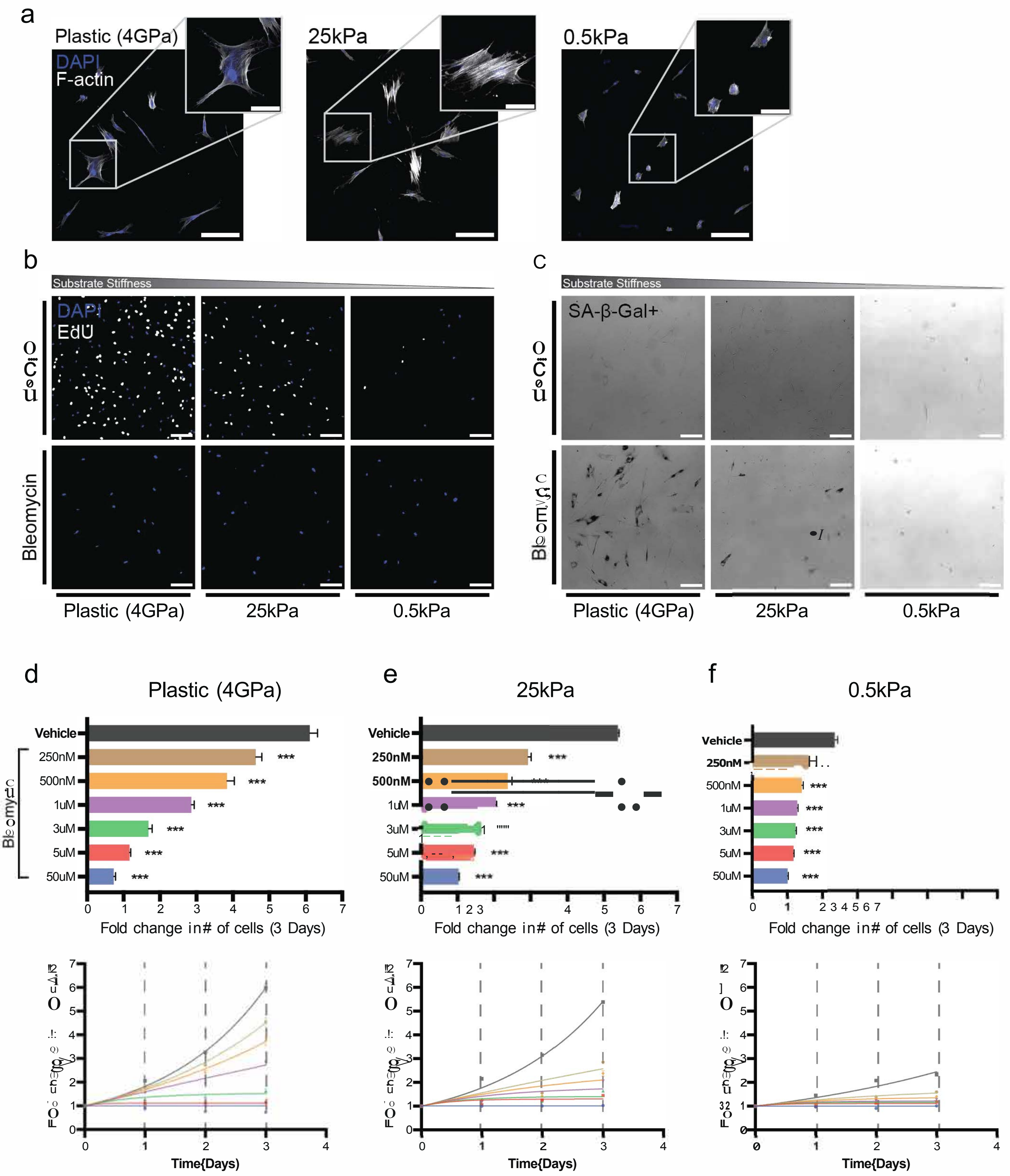

**Supplemental Figure 1 – Bleomycin induction inhibits GT22 fibroblast proliferation on Plastic (4GPa), 25 kPa, and 0.5 kPa substrates** (**a**) Representative Images of non-senescent GT22 primary dermal fibroblasts on Plastic, 25kPa, and 0.5kPa collagen coated subtrates. Primary scalebar 200µm, inlayed scalebar 75µm. (**b**) EDU staining 7 days post senescence induction. White cells indicate EDU+ proliferating cells, blue nuclei, scalebar 200µm (**c**) Representative SA-B-gal staining for GT22 fibroblasts 7 days after induction on Plastic, 25 kPa and 0.5 kPa substrates show SA-B-gal not clearly expressed at soft substrate conditions, scalebar 200µm. (**d**) WI-38 fibroblast proliferation response. Bleomycin 3-day endpoint bar plots and corresponding dose response curves showing fold change in number of fibroblasts vs time for cells subjected to Vehicle, 250nM, 500nM, 1µM, 3µM, 5µM, 50µM: (**d**) Plastic (4GPa), (**e**) 25 kPa, and (**f**) 0.5 kPa matrices. Error bars represent SEM of technical replicates, statistics evaluated via one-way ANOVA, with Tukey multiple comparison test to vehicle control; data presented are from 1 biological replicate with each having 5-10 technical replicates. (***P <0.001, **P < 0.01, *P < 0.05).

Supplemental Figure 2

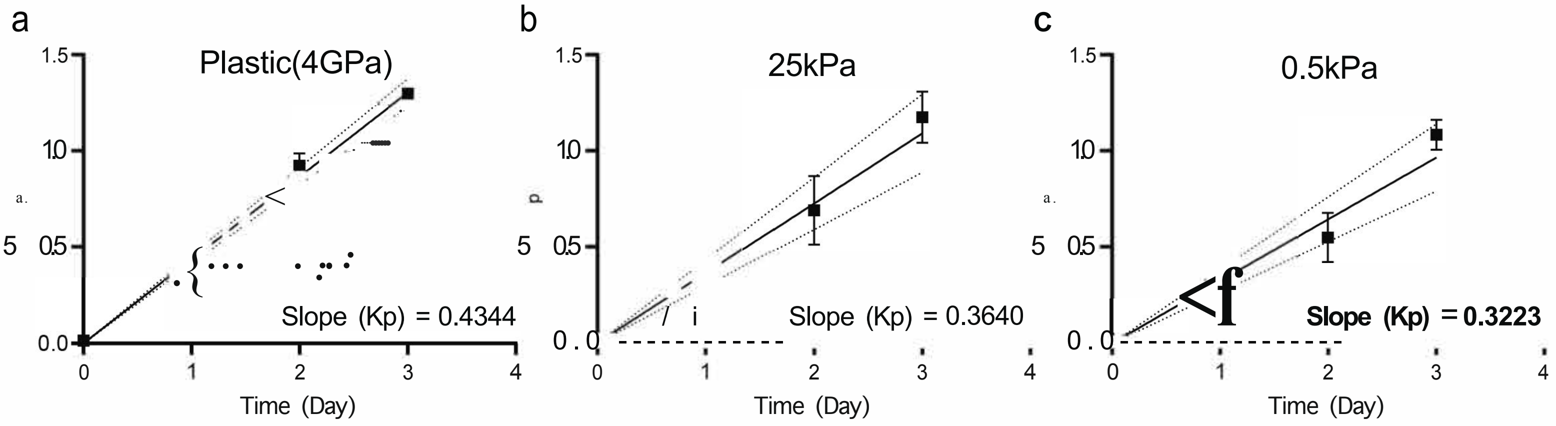


d

Ks Summary Table

| Condition | Plastic | | 25kPa | | 0.5kPa | |
|---|---|---|---|---|---|---|
| | Mean | R2 | Mean | R2 | Mean | R2 |
| Vehicle | 0 | 0.92 | 0 | 0.58 | 0 | 0.65 |
| 250nM | 0.20 | 0.62 | 0.89 | 0.21 | 1.26 | 0.23 |
| 500nM | 0.32 | 0.56 | 1.24 | 0.15 | 1.88 | 0.12 |
| 1uM | 0.54 | 0.61 | 1.72 | 0.22 | 3.73 | 0.03 |
| 3uM | 1.00 | 0.33 | 5 | * | 5 | * |
| 5uM | 5 | * | 5 | * | 5 | * |
| 50uM | 5 | * | 5 | * | 5 | * |

*Model hit constraint

**Supplemental Figure 2 – Unconstrained growth constants for cells cultured on Plastic, 25 kPa, and 0.5 kPa substrates**. (**a**) Log transformed unconstrained cell proliferation for Plastic (4GPa) vs time. Linear regression fit to extract drug free growth rate. (**b**) 25 kPa unconstrained growth rate, (**c**) 0.5 kPa unconstrained growth rate. (**e**) Summary table of Ks fits. (**f**) Model fit to N biological = 3, n technical = 3-5.

Supplemental Figure 3

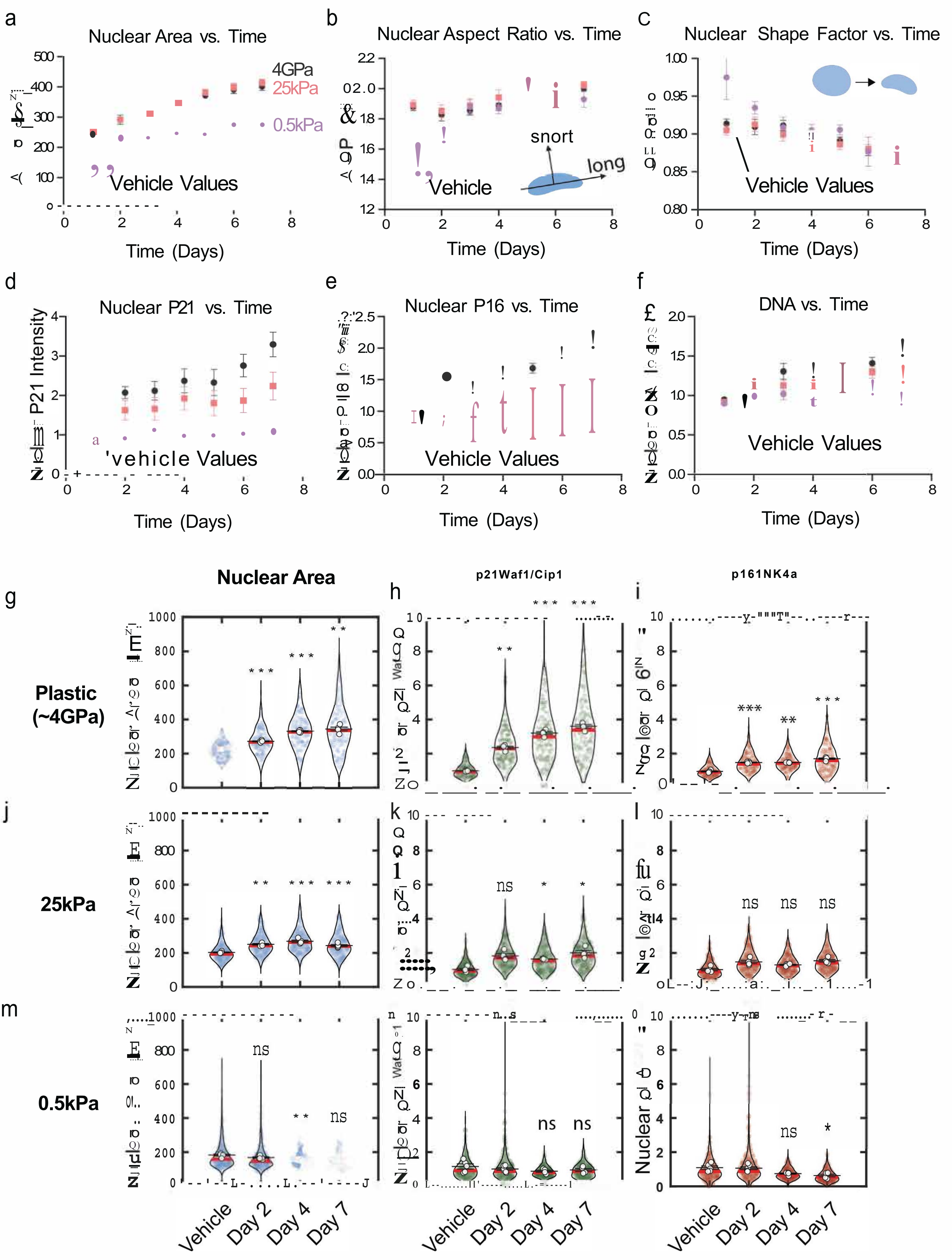

**Supplemental Figure 3 – Evolution of morphological and protein phenotypes in senescence.** (**a**) Nuclear area of senescent GT22s on 0.5 kPa, 25 kPa and 4GPa (Plastic) substrates. (**b**) Nuclear aspect ratio increased in senescence cells. (**c**) nuclear shape factor decreases to less rounded, more tortuous morphology. (**d**) Nuclear $p21^{WAF1/CIP1}$ expression normalized to DPBS control increases in senescent cells on 25 kPa and Plastic substrates. (**e**) Nuclear $p16^{INK4a}$ normalized to PBS Control increases slowly increases in 25 kPa and Plastic substrates but remains suppressed on soft substrates. (**f**) DAPI intensity vs time shows reduced DNA content for soft senescent fibroblasts with time. (**g-o**) Wi-38 senescent cells experiencing a 4GPa plastic substrate exhibit an increased (**g**) nuclear area ($\mu m^2$), (**h**) nuclear expression of $p21^{WAF1/CIP1}$ and (**i**) nuclear expression of $p16^{INK4a}$ with time. WI-38 fibroblasts on 25kPa substrates show increased (**j**) nuclear area but reduced (**k**) nuclear $p21^{WAF1/CIP1}$ and (**l**) nuclear $p16^{INK4a}$ with time. 0.5kPa fibroblasts show (**m**) decreased nuclear growth and negligible expression of (**n**) nuclear $p21^{WAF1/CIP1}$ and (**o**) nuclear $p16^{INK4a}$ with time. For GT22 immunofluorescence data error bars represent SEM of biological variation. For WI-38 IF data, error bars are SEM of technical replicates. Single cell IF statistics evaluated via one-way ANOVA, with Tukey multiple comparison test to vehicle control. GT22: N=3, n = 3-5, WI-38: N = 1, n = 3-5 (***P <0.001, **P < 0.01, *P < 0.05).

### Supplemental Figure 4

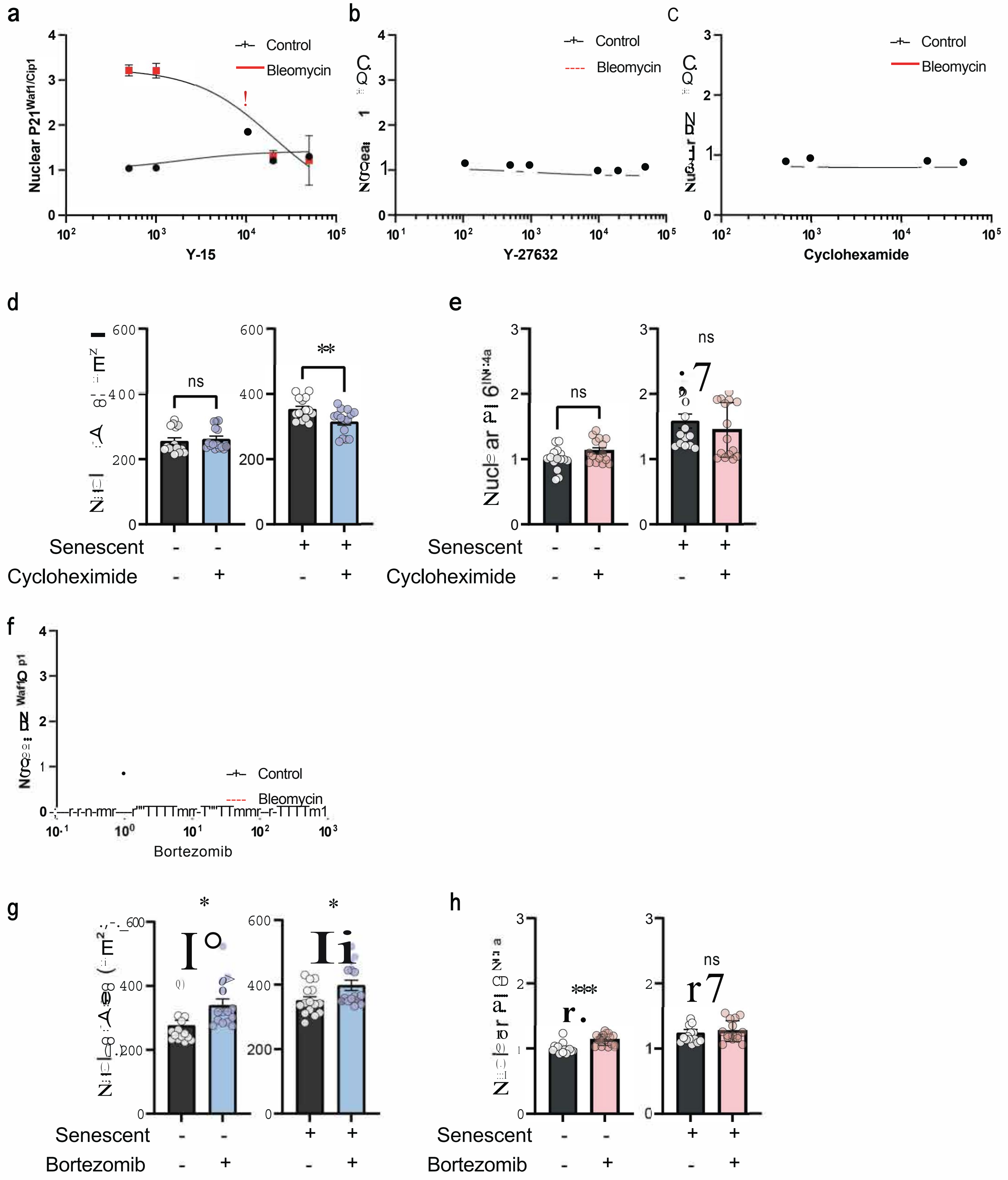

**Supplemental Figure 4 – Small molecule intervention in the senescence phenotype**. (**a**) dose response for FAKi (Y-15) vs nuclear $p21^{WAF1/CIP1}$, (50 μM, 20 μM, 10μM, 1 μM, 500nM), red curve shows bleomycin treated senescent fibroblasts, black vehicle control. (**b**) dose response for ROCKi (Y-27632) vs nuclear $p21^{WAF1/CIP1}$, (50 μM, 20 μM, 10μM, 1 μM, 500nM, 100nM). (**c**) dose response for Cycloheximide vs nuclear $p21^{WAF1/CIP1}$, (50 μM, 20 μM, 1 μM, 500nM). Early senescent fibroblasts treated 1 μM Cycloheximide showed limited changes to (**d**) nuclear area and (**e**) nuclear $p16^{INK4a}$. (**f**) dose response for Borezomib vs nuclear $p21^{WAF1/CIP1}$, (100nM, 10nM, 1nM, 500pM). Intervention in ubiquitin mediated protein degradation using 100nM Bortezomib increased (h) nuclear area with limited changes in (**g**) nuclear $p16^{INK4a}$. Error bars represent SEM of biological variation, N biological = 3, n technical = 3-5. (***P <0.001, **P < 0.01, *P < 0.05). Statistics evaluated using Welch's t-test.
